# A chaotrope-based approach for rapid *in vitro* assembly and loading of bacterial microcompartment shells

**DOI:** 10.1101/2024.10.31.621445

**Authors:** Kyleigh L. Range, Timothy K. Chiang, Arinita Pramanik, Joel F. Landa, Samuel N. Snyder, Xiaobing Zuo, David M. Tiede, Lisa M. Utschig, Eric L. Hegg, Markus Sutter, Cheryl A. Kerfeld, Corie Y. Ralston

## Abstract

Bacterial microcompartments (BMCs) are proteinaceous organelles that self-assemble into selectively permeable shells that encapsulate enzymatic cargo. BMCs enhance catalytic pathways by reducing crosstalk among metabolites, preventing harmful intermediates from leaking into the cytosol, and increasing reaction efficiency via enzyme colocalization. The intrinsic properties of BMCs make them attractive for biotechnological engineering. However, *in vivo* expression methods for shell synthesis have significant drawbacks that limit the potential design space for these nanocompartments. Here we describe the development of a new, efficient, and rapid method for *in vitro* assembly of BMC shells from their protein building blocks. Our method enables large-scale construction of BMC shells by utilizing urea as a chaotropic agent to control self-assembly, and provides an approach for encapsulation of both biotic and abiotic cargo under a broad range of reaction conditions. We demonstrate an enhanced level of control over the assembly of BMC shells *in vitro* and expand the design parameter space for engineering BMC systems with specialized and enhanced catalytic properties.

---

Membrane-bound organelles were once thought to be an exclusive characteristic of eukaryotes, but partitioned structures in bacteria also provide a wide diversity of subcellular organizational systems in organisms across domains. In contrast to lipid membrane-bound eukaryotic organelles, bacterial microcompartments (BMCs) use only proteins^1,2^ to encapsulate various metabolic pathways.^3^ The carboxysome, for example, is a BMC that is found among cyanobacteria and other autotrophic bacteria that enhances carbon fixation through colocalization of the enzyme RuBisCO (ribulose-1,5-biphosphate carboxylase/oxygenase) and concentrating its substrate, CO_2_, thereby overcoming some of RuBisCO’s inefficiencies.^4,5^ By encapsulating catalysts within a selectively permeable protein membrane, BMC shells increase enzyme efficiency through colocalization, prevent metabolic crosstalk between competing substrates, and eliminate the spillage of toxic or volatile intermediates into the cytosol.^3,6^ The polyhedral BMC shell is composed of hexameric, pseudo-hexameric, and pentameric “tiles” that are evolutionarily conserved across different taxa.^1^ In BMCs, the most abundant shell tiles, hexamers, consist of six identical protomers containing a single pfam00936 domain (BMC-H proteins). BMC-T proteins, a fusion of two copies of the pfam00936 domain, form trimers that resemble the hexamers in size and shape and are a less abundant component of shell facets.^1,2^ Pentamers (BMC-P proteins) that cap the vertices of the polyhedral shell are composed of five protomers of the pfam03319 domain.^7^

Since the first report of a recombinantly expressed BMC system,^8^ purification and characterization of native BMCs^9–12^ and empty shells^13–18^ has contributed to increased understanding of shell structure, function, and assembly. Although its native function is unknown, the microcompartment from *Haliangium ochraceum* (HO) provides robust *in vivo* shell assembly through recombinant expression of its shell protein constituents.^19^ Through manipulation of a synthetic operon, three main shell types (full, minimal, and minimal wiffle shells) can be purified from heterologous expression.^20^ Full HO shells incorporate three distinct trimer proteins of which two (BMC-T2 and BMC-T3) form dimers that are double-layered. Minimal (HTP) and minimal wiffle (HT) HO shells both contain only a single type of trimer (BMC-T1, referred to here onward as BMC-T) with BMC-H arranged in a T=9 icosahedra, differing in that minimal wiffle shells lack pentamers at the 12 icosahedral vertices. The stoichiometric ratio of shell tiles in a minimal HTP shell is 60:20:12 BMC-H:BMC-T:BMC-P, or 60:20 BMC-H:BMC-T in a minimal wiffle HT shell.^21^

*In vivo* expression and purification of BMC shells is time and labor intensive and does not offer precise control of composition in self-assembly. Moreover, shells expressed *in vivo* adventitiously capture unwanted contaminants from the cytosolic milieu.^22^ Previously, an *in vitro* assembly (IVA) method for constructing BMC shells was reported that overcame several limitations of recombinantly-expressed shells, offering a higher degree of control over shape, size, and cargo content.^23^ This method relied on enzymatic activity to cleave a genetically introduced blocking group on the BMC-H to initiate shell assembly; this inherently limits reaction conditions to the narrow range that is conducive to the enzymatic function. We have developed a chaotrope-based method for *in vitro* assembly of BMC shells. This method provides a powerful new tool for bioengineering efforts as it allows more precise control of shell composition, significantly increases both speed and efficiency, and expands the range of reaction conditions for assembly. Furthermore, it provides increased flexibility in the choice of BMC tile building blocks enabling a broader range of shell functionalization, thus broadening the spectrum for controlled catalysis within these partitioned systems.

## Results

### *In vitro* assembly of BMC shells from their constituent protein tiles

We prepared purified shell proteins and combined them according to the reaction schemes shown in **Figure 1**. Complete sequences of all proteins used in this study are listed in **Table S1**, and their molecular weights are listed in **Table S2**. BMC-T and BMC-P tiles were heterologously expressed in *E. coli* and purified using affinity chromatography, and were either dialyzed into Tris buffer or subjected to size exclusion chromatography (SEC). When HO BMC-H is heterologously expressed in *E. coli* in the absence of other shell proteins, it forms inclusion bodies that can be purified as a suspension of BMC-H sheets^24,25^ (**Figure 1A**). These supramolecular sheet structures are insoluble, and the constituent BMC-H tiles are therefore not viable for assembling BMC shells. We obtained assembly-competent BMC-H tiles in soluble form by disassembling sheets with 500 mM urea (**Figure 1A**). We found that this concentration of urea is high enough to disrupt BMC-H–BMC-H interactions within sheets without inhibiting BMC-H–BMC-T and BMC-H–BMC-P tile interfaces necessary for shell assembly. The solubilized BMC-H, BMC-T, and BMC-P tiles were pure and monodisperse as determined by dynamic light scattering (DLS) (**Figure S1**).

**Figure 1.**
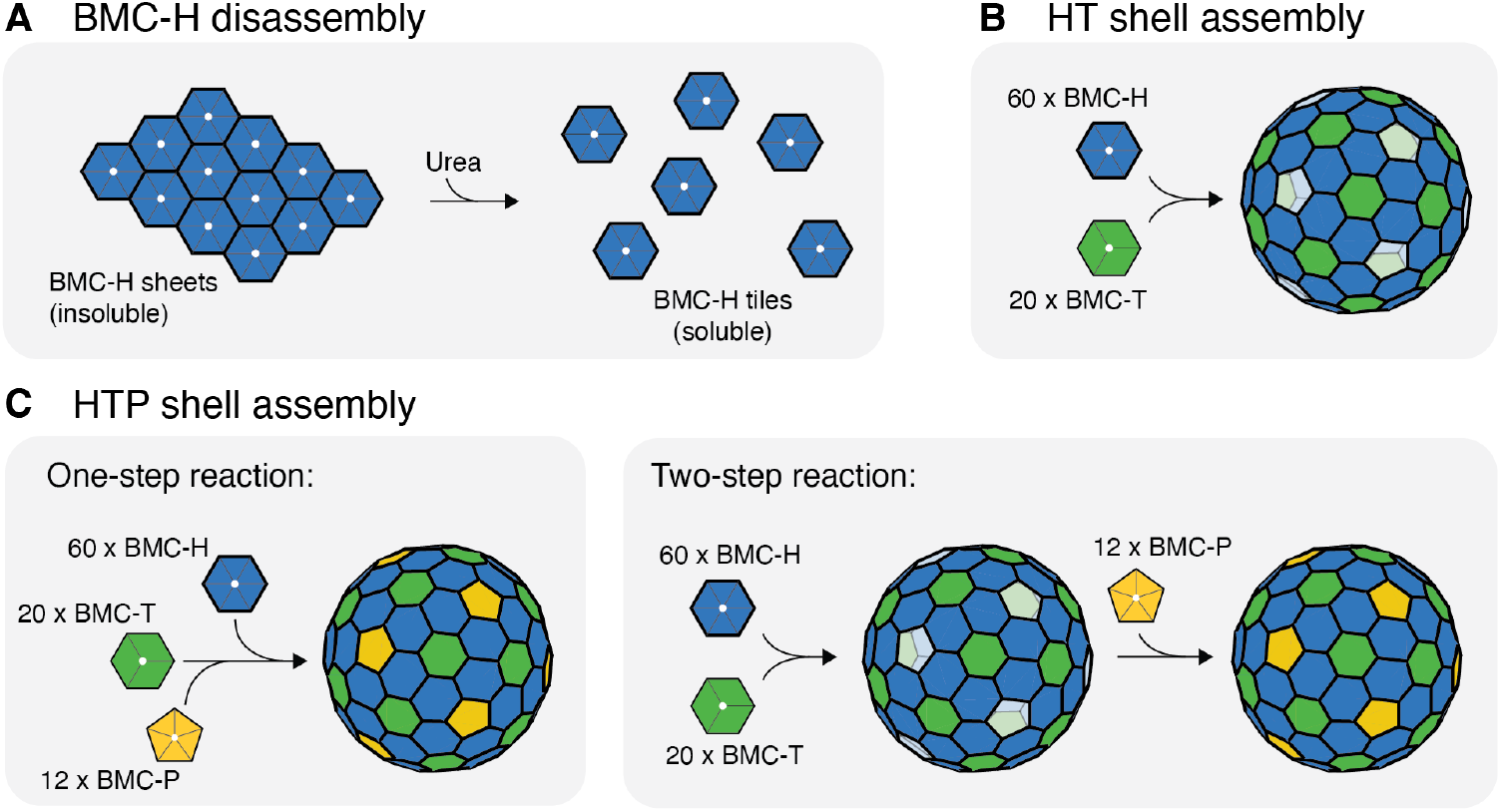
Schematic representation of the chaotrope-based approach to *in vitro* BMC shell assembly. **A**. BMC-H hexamers form sheets that disassemble into assembly-competent tiles upon the addition of urea. **B**. Combining solutions of heterologously expressed and purified BMC-H and BMC-T shell proteins results in the *in vitro* assembly of HT shells. **C**. Combining solutions of all three BMC shell proteins results in the *in vitro* assembly of HTP shells. HTP shells can be assembled upon mixing via a one-step (BMC-H + BMC-T + BMC-P) or two-step (BMC-H + BMC-T first, then BMC-P) addition.

BMC shells were assembled by mixing purified solutions of BMC tiles according to the assembly reaction schemes shown in **Figure 1B** and **Figure 1C**. To assemble HT or HTP shells, we combined BMC-T and BMC-P in Tris buffer before adding BMC-H with vigorous mixing, followed by incubation at 4°C for 24 hours, mirroring the protocol from Hagen *et al*.^23^ We also tested assembly after 2 minutes and 1 hour to observe how assembly duration affects reaction efficiency. The HT shell assembly reactions were performed at 1 mg/mL BMC-H and 0.33 mg/mL BMC-T, following the expected stoichiometric molar ratio for a HT wiffle shell (3:1 BMC-H:BMC-T). The addition of pentamers in the HTP shell assembly reaction fills the 12 vacancies of HT shells vertices and creates a microenvironment within the shell that is essentially sealed from entry and exit of large solutes. We assembled HTP shells using a one-step or two-step protocol (**Figure 1C**). For the one-step HTP assemblies, we combined all three shell proteins in the IVA reaction with BMC-P in a five-fold excess; 0.33 mg/mL BMC-T and 1 mg/mL BMC-P tiles were mixed into assembly buffer before the addition of 1 mg/mL BMC-H. We found that BMC-H will initiate assembly with BMC-T; therefore in order not to dilute urea or initiate shell assembly, it was the last shell protein to be added to the reaction buffer. The reaction was similarly left to incubate at 4°C. For the two-step assembly, we first generated minimal wiffle shells that were then capped with BMC-P, a method previously shown successful for *in vivo* generated HO shells.^20^ We used the same conditions as in a recent report by Snyder *et al*.^26^ and assembled verified HT shells before adding a five-fold excess of BMC-P for a 30 minute incubation period.

### Characterization of assembled shell components and structure

To verify the formation of BMC shells, we analyzed the components and structure of the assemblies using SDS-PAGE, DLS, negative stain transmission electron microscopy (TEM), and small angle X-ray scattering (SAXS). The assembled shells were purified using SEC, separating them from individual tiles and removing residual urea (**Figure 2A–C**). In all assembly reactions, most of the protein eluted in the column void volume, indicating the presence of large assemblies. The size of the species in these fractions were measured to be approximately 40 nm with low polydispersity, as determined by DLS (**Figure S1**), which is consistent with previously reported HO BMC shell diameters.^19,21,27^ Direct imaging using negative stain TEM analysis of the shell fractions provided confirmation of assembled shell structures, and SDS-PAGE analysis verified the composition of the shells (**Figure 2D**). Shell assemblies were also characterized with SAXS to verify assembly size and structure. The SAXS profiles depict oscillatory features that reflect the core-shell particle structures (**Figure 2E**). The attenuation of the oscillations in the experimental data, compared to the model SAXS spectrum, is due in part to the polyhedral structure of the shell that deviates from an ideal sphere. In comparison with HTP shells assembled *in vivo* (**Figure S2**), the SAXS profiles of the *in vitro* assembled HTP shells exhibit slightly more attenuation of the oscillatory features and a shift in the first minima/maxima oscillation to a lower Q value, suggesting a broader size distribution in the *in vitro* assembled sample. Nonetheless, comparison of the *in vitro* assembled HT and HTP shells with *in vivo* expressed HTP shells and modeled hollow core-shell spheres illustrates consistency across all structures.

**Figure 2.**
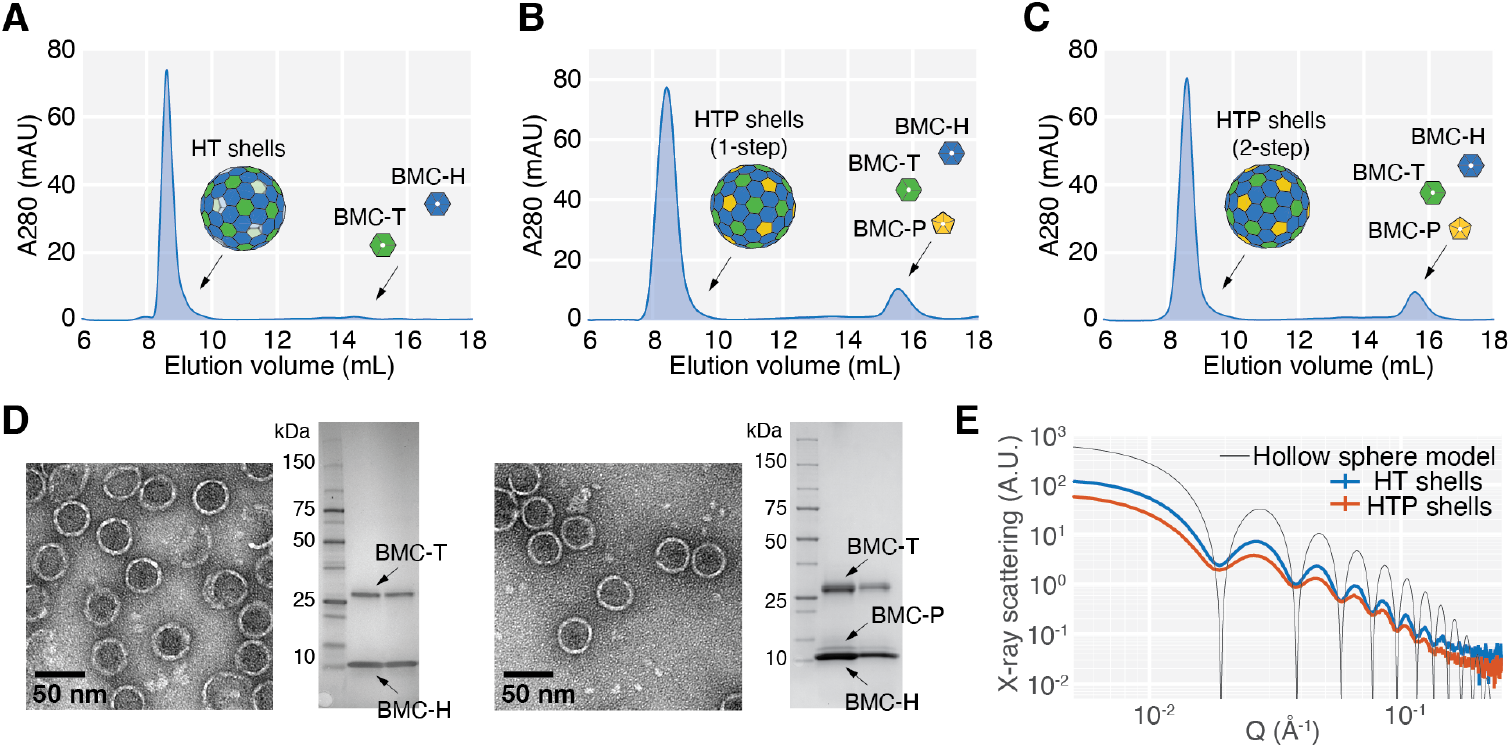
Characterization of *in vitro* assembled HT and HTP shells. **A**. SEC chromatogram for *in vitro* assembled HT shells. The main elution peak contains monodisperse 40 nm hollow spheres made from BMC-H and BMC-T. **B–C**. SEC chromatograms for *in vitro* HTP shells assembled via 1-step (**B**) and 2-step (**C**) reactions, with a five-fold stoichiometric excess amount of BMC-P. The main elution peak contains monodisperse 40 nm hollow spheres made from BMC-H, BMC-T, and BMC-P. **D**. Negative stain TEM micrographs and SDS-PAGE analyses of HT shells (left) and 1-step HTP shells (right). **E**. SAXS spectra for *in vitro* assembled HT and HTP shells and simulated spectrum for a hollow sphere model with inner radius of 150.5 Å and shell thickness of 25 Å.

### *In vitro* shell assembly reaction efficiency and speed

We next sought to quantify the efficiency and speed of the assembly. We define assembly efficiency as the amount of protein in shell form as a fraction of the total amount of protein in both shell form and in unassembled tiles. We determined the amount of protein from SEC fractions by integrating the UV 280 absorbance peaks. Because the scattering contributions from the large assembled shells are non-negligible, these measurements were also compared to the results from a BCA assay (**Figure S3**). In assemblies that utilize the expected stoichiometry of tiles, we find that assembly efficiency reaches between 75–94% (**Figure 3A** and **Figure S3**). This is significantly higher than that from the previous IVA method,^23^ which reported an approximate efficiency of 20%. Moreover, since the previous IVA method relied on an enzymatic cleavage step, maximum assembly efficiency was achieved only for overnight incubations. Likewise, the overall cost of our method is lower since it does not require either a separately purified or commercially obtained protease.

**Figure 3.**
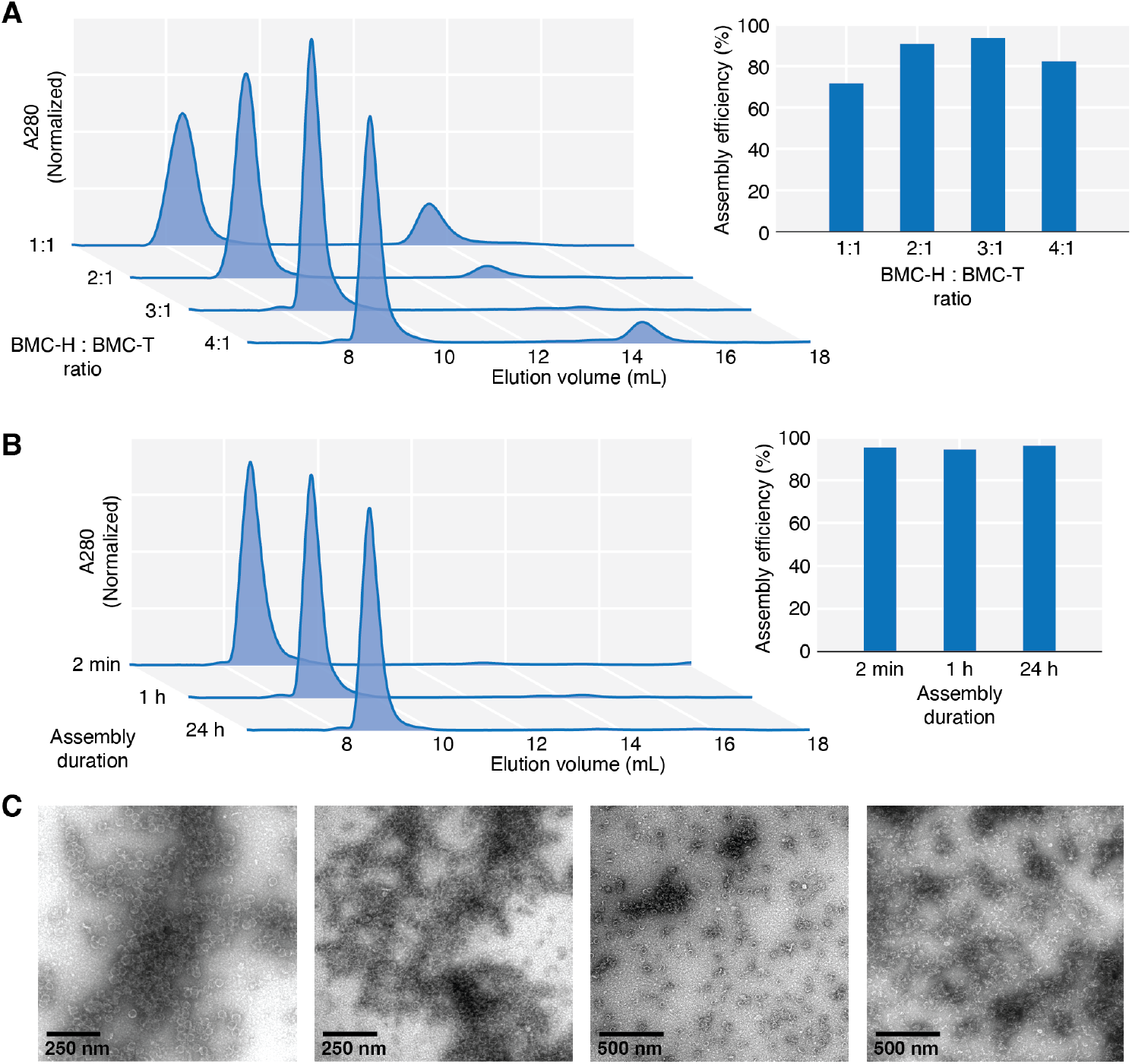
*In vitro* assembly of HT shells is efficient and fast. **A**. SEC chromatograms of HT shell assembly reactions containing varying BMC-H:BMC-T stoichiometries. The bar graph shows that the assembly efficiency is highest at the expected stoichiometric ratio of 3:1 BMC-H to BMC-T. Efficiency is defined as the area of the chromatogram peak containing shells divided by the total area. **B**. SEC chromatograms of HT shells assembled with varying duration at a fixed BMC-H:BMC-T ratio of 3:1. The bar graph shows that the reaction efficiencies among 2-minute, 1-hour, and 24-hour assemblies are indistinguishable, suggesting that HT shell assembly is complete in under 2 minutes. **C**. Representative TEM micrographs of shells assembled directly on carbon grids and stained immediately.

We sought to minimize the time between assembly and purification to mitigate potential adverse effects on biotic cargo due to prolonged urea exposure. We tested the completion of the assembly reaction after 2 minutes, 1 hour, and 24 hours. In all cases, only assembled shells elute from the size exclusion column (**Figure 3B**). Additionally, we performed an assembly reaction and immediately applied it on grids for negative staining TEM analysis, essentially capturing the state of the assembly reaction after 10 seconds (**Figure 3C**). TEM images showed many assembled shells, suggesting that individual shells assemble on timescales much faster than our ability to detect them with this method.

### Loading of non-native, catalytically active cargo

#### Targeted loading of the enzyme NrfA via shell protein conjugation

To demonstrate the application of *in vitro* BMC shell assembly for the encapsulation of catalytic cargo, we investigated the targeted loading of a non-native enzyme, cytochrome *c* nitrite reductase (NrfA) from *Geobacter lovleyi*.^28^ Using the SpyTag-SpyCatcher system,^29,30^ we covalently tethered cargo enzyme molecules to modified shell proteins. Specifically, we expressed and purified a construct of the BMC-T subunit with a SpyTag linker cloned to a loop region of the protein, previously shown to tolerate insertions.^20^ The linker is located on the interior facet of the shell component in its assembled state. We investigated the ability of the BMC-T tile modified with SpyTag (referred to here onward as _SpyT_BMC-T) to assemble into HT and HTP shells *in vitro* and found that they were successfully incorporated into assembled shells, though at a slightly lower efficiency compared to untagged BMC-T tiles (**Figure S4**). We also cloned, expressed, and purified NrfA with a SpyCatcher linker attached to the N-terminus (referred to here onward as NrfA_SpyC_).

Because BMC-T tiles consist of three identical protomer subunits, the _SpyT_BMC-T construct results in a tile that contains three SpyTags. Binding all three SpyTags with NrfA_SpyC_ makes the protein considerably more bulky and less viable for assembly. To prevent oversaturation of single _SpyT_BMC-T tiles, we conjugated using an approximate ratio of 6 tiles per 1 enzyme or 3 enzymes per one shell. We conducted the conjugation reaction before assembly for 1 hour on ice. Following incubation, we diluted the mixture of linked _SpyT_BMC-T and NrfA_SpyC_ in assembly buffer before the addition of BMC-H. We incubated the assembly reaction for 1 hour on ice before loading the sample onto a size exclusion column. NrfA contains five internal c-type hemes absorbing at 410 nm in the Fe^3+^ oxidation state, allowing us to track the elution of the enzyme at a wavelength that is distinct from shell protein absorption (**Figure 4A**).

**Figure 4:**
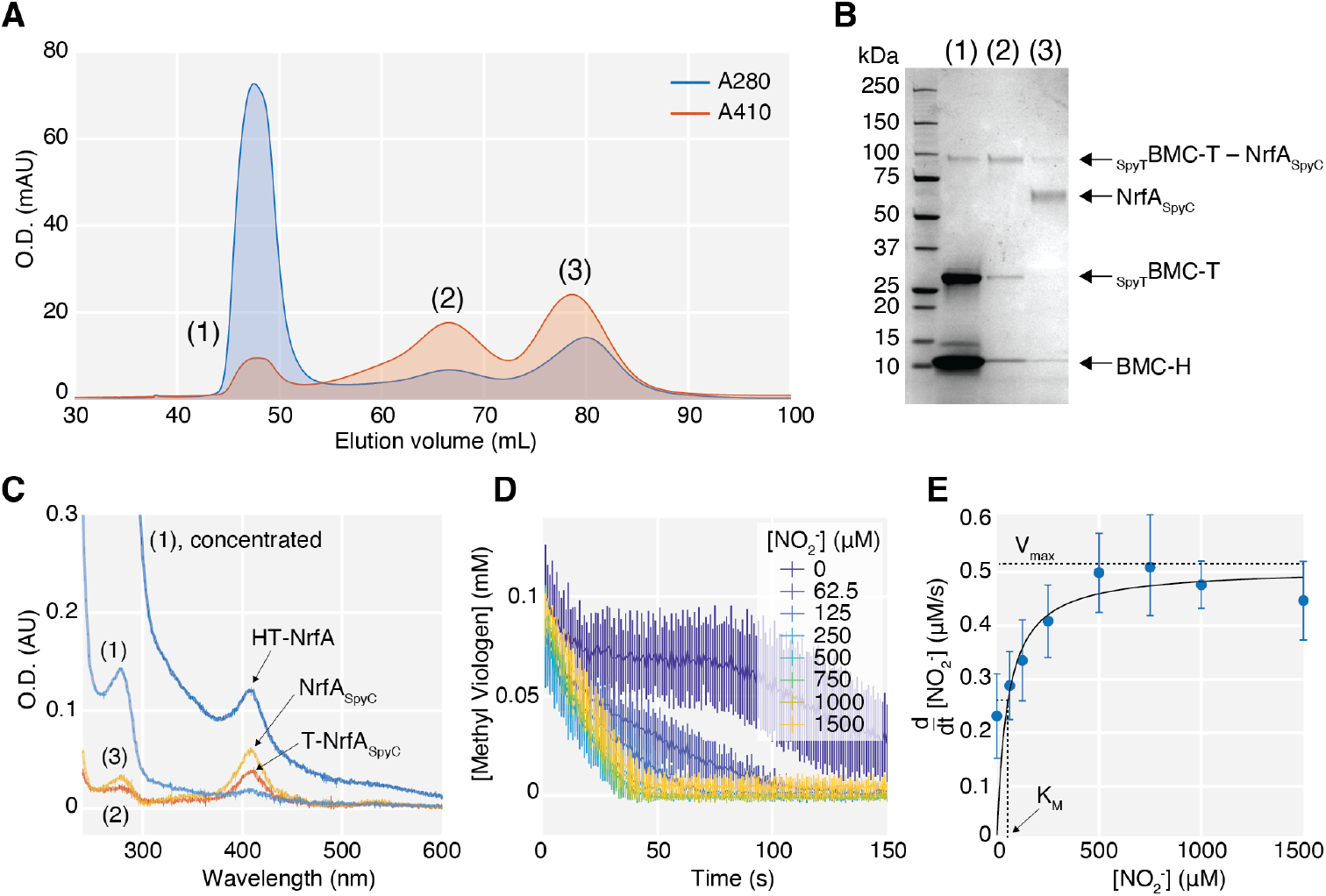
*In vitro* encapsulated NrfA_SpyC_ is catalytically active. **A**. SEC chromatogram showing purification of NrfA_SpyC_-loaded HT wiffle shells from unincorporated BMC-H, _SpyT_BMC-T, and NrfA_SpyC_. **B**. SDS-PAGE analysis shows _SpyT_BMC-T–NrfA_SpyC_ conjugate in the elution peak (1) fraction containing HT shells, and unincorporated conjugated _SpyT_BMC-T–NrfA_SpyC_ and unconjugated NrfA_SpyC_ in peaks (2) and (3), respectively. **C**. UV-Vis spectra of SEC fractions for peaks (1)–(3), and a concentrated solution of (1). **D**. Activity assay of NrfA_SpyC_ inside HT shells in peak (1). Kinetic traces of the concentration of reduced methyl viologen at various concentrations of nitrite show the rates of substrate turnover. **E**. Michaelis-Menten curve, with V_max_ = 0.502 ± 0.040 μM/s and K_M_ = 49.45 ± 25.47 μM.

We observed conjugated _SpyT_BMC-T–NrfA_SpyC_ in the same fractions as assembled shells as verified by SDS-PAGE (**Figure 4B**) and confirmed that conjugation of enzyme to tile did not disrupt or change shell morphology using DLS and TEM (**Figure S5**). Control SEC runs without the addition of BMC-H showed that the _SpyT_BMC-T–NrfA_SpyC_ conjugate as well as unconjugated _SpyT_BMC-T and NrfA_SpyC_ did not elute in the void volume (**Figure S6**). The concentration of the enzyme inside the shells was measured via UV-Vis absorbance of the heme Soret peak at 410 nm (**Figure 4C** and **Figure S7**) and was found to be approximately 112 nM. Using the absorbance at 280 nm to extract an estimation for the concentration of HT shells, we calculated yields of roughly 425 nM shells, indicating an approximate loading efficiency of 26%, or roughly 1 enzyme per 4 HT shells. The large amount of unconjugated NrfA_SpyC_ (peak (3) in **Figure 4A–C**) suggests that the encapsulation efficiency can be improved by changing reaction parameters, such as increasing the duration of incubation of the enzyme with _SpyT_BMC-T before the initiation of assembly upon addition of BMC-H.

To evaluate the catalytic activity of the encapsulated enzyme, we performed a Michaelis-Menten enzyme activity assay on the SEC-purified HT shells loaded with NrfA_SpyC_ at a target concentration of 1 nM NrfA_SpyC_ in the final reaction mixture. NrfA catalyzes the conversion of nitrite (NO_2_^-^) to ammonium, driven by electron donation from dithionite-reduced methyl viologen. To measure the rate of nitrite reduction by NrfA_SpyC_, we monitored the change in absorbance, as the solution containing reduced methyl viologen turns from blue to clear upon oxidation by the turnover of nitrite by NrfA_SpyC_ (**Figure 5D**). The rate of reaction at early time points and at various substrate concentrations were fit to a hyperbolic Michaelis-Menten curve, from which *K*_M_ was measured to be 49.45 ± 25.47 μM, and *V*_max_ was 0.502 ± 0.040 μM/s (**Figure 5E**). The *K*_M_ is higher than that of the wild-type enzyme free in solution (27 ± 2 μM),^28^ suggesting a decrease in accessibility of the substrate to the active site. This is potentially due to the barrier imposed by the HT shell. The rate of catalysis, *k*_cat_, was inferred to be about 536 ± 68 μmol NO_2_^-^ min^-1^ mg^-1^ enzyme. While this value of *k*_cat_ is lower than that of the wild-type enzyme (1291 ± 34 μmol NO_2_^-^ min^-1^ mg^-1^ enzyme),^28^ measurements of these parameters are in general sensitive to slight differences in experimental conditions and additionally our enzyme concentration determination has a large margin of error. Importantly, these data show that the NrfA_SpyC_ inside HT shells remain catalytically active after conjugation and encapsulation.

**Figure 5.**
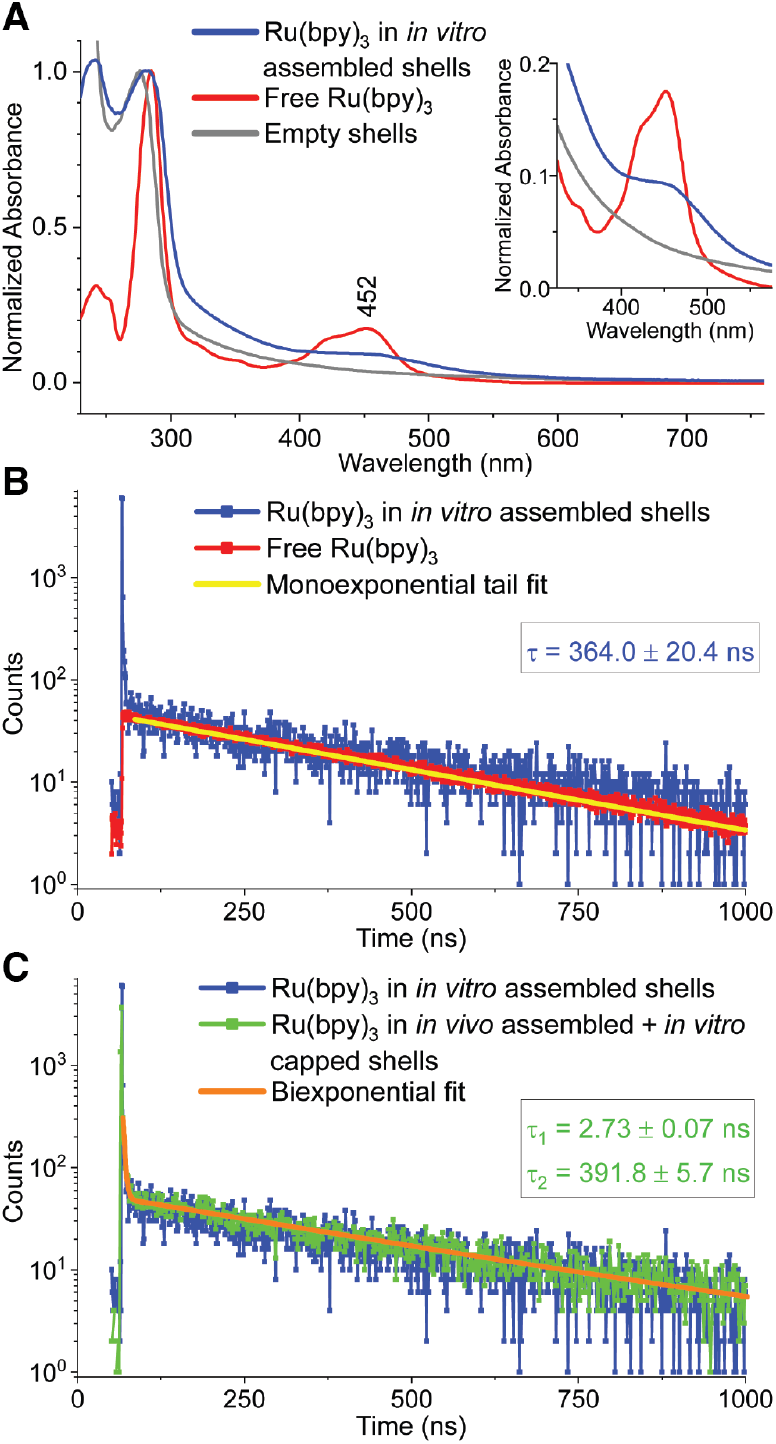
Spectroscopic data of the abiotic cargo molecule, Ru(bpy)_3_, encapsulated in HTP shells by the one-step *in vitro* assembly method. **A**. UV-Vis spectra. Data are normalized by absorption at the bands around 277–285 nm in the UV-region and a magnification is shown for the region containing MLCT bands (inset). **B–C**. TCSPC photoluminescence data are shown with indicated lifetimes derived from fitting with **B**. monoexponential or **C**. biexponential functions. Emission was measured at 620 nm using pulsed laser excitation at 445.8 nm.

#### Encapsulation of Ru(bpy)_3_ photosensitizer via passive diffusion

To demonstrate an application of our *in vitro* BMC shell assembly for the encapsulation of a different non-native cargo, we investigated loading of abiotic cargo molecules. A recent study showed that the abiotic molecule and benchmark photosensitizer, [Ru(bpy)_3_]^2+^, could be encapsulated via diffusion through the vacancies in *in vivo* assembled HT shells and then trapped in the shell lumen upon *in vitro* capping with pentamer to form sealed HTP shells^26^. Our one-step *in vitro* assembly method outlined here was able to successfully encapsulate Ru(bpy)_3_ at comparable cargo loading efficiencies to the former *in vivo* assembly *+ in vitro* capping method (**Table S3**). Ru(bpy)_3_Cl_2_•6H_2_O was dissolved in the shell protein solutions to a concentration of 13 mM immediately prior to initiation of shell assembly and subsequent separation of unassembled proteins and excess, unencapsulated Ru(bpy)_3_ (see **Methods and Materials** in **Supporting Information**). This *in vitro* assembly resulted in cargo loading of ∼20–22 Ru(bpy)_3_ molecules per shell and required no specific interactions between shell proteins and abiotic cargo molecules for successful encapsulation. It should also be noted that these data indicate that the *in vitro* assembly method results in a significant population of sealed, complete shells because control studies have shown that Ru(bpy)_3_ molecules will not remain in the shell lumen unless the shell is complete, *i.e*. without any vacancies from a missing shell protein.^26^

UV-Vis characterization of the *in vitro* assembled shells harboring Ru(bpy)_3_ showed a metal-to-ligand charge transfer (MLCT) band centered around 452 nm as expected from the spectrum of free Ru(bpy)_3_, although the shape of the band is somewhat skewed by scattering from the shell that raises the baseline (**Figure 5A**). The excited state lifetime was evaluated with time-correlated single photon counting (TCSPC) photoluminescence spectroscopy. The spectrum of Ru(bpy)_3_ encapsulated within the *in vitro* assembled shells has two time components, one of which is in agreement with that reported in literature and measured here for free Ru(bpy)_3_, with a lifetime of ∼360 nanoseconds when tail-fitting the data with a monoexponential function (**Figure 5B**).^31,32^ There is a much faster decaying species that is absent from the free Ru(bpy)_3_ spectrum, but overlays well with the spectrum of Ru(bpy)_3_ in shells assembled by the former combined *in vivo* assembly + *in vitro* capping method with a determined lifetime of ∼3 ns when fit with a biexponential function (**Figure 5C**). This faster decaying time component was attributed to possible intermolecular interactions of the Ru(bpy)_3_ excited state with neighboring excited or ground state Ru(bpy)_3_ molecules,^26^ drawing upon a literature precedent for Ru(bpy)_3_ in confinement in zeolites.^33^ Collectively, the spectroscopic data indicate that the photophysical properties of Ru(bpy)_3_ are maintained after being encapsulated by shells via the *in vitro* assembly method and bode well for future applications of Ru(bpy)_3_ and other compounds, biotic or abiotic, in confinement within BMC architectures.

## Discussion

Our chaotropic *in vitro* assembly method provides several significant advancements in the synthesis of bacterial microcompartment shells. Among the most notable of these improvements are the efficiency and speed of shell assembly. In contrast to the *in vitro* assembly method utilizing enzymatic cleavage, our method more than triples efficiency; nearly all constituents are used up to form monodisperse shells. By combining assembly competent BMC-H and BMC-T tiles, our method drastically reduces reaction incubation time to under 2 minutes. The robustness of the method in a variety of conditions is validated by assemblies successfully performed in a range of urea concentrations (100 mM and 900 mM) as shown by SAXS (**Figure S2**). Furthermore, in these higher urea solutions the SAXS profiles indicate lower heterogeneity (**Figure S2**), suggesting that it might be possible to control shell size distribution with urea concentration. Furthermore, this method is able to generate the highest purity of shell samples through our ability to assemble shells using only their constituent protein building blocks. It diminishes the presence of contaminating cytosolic proteins or enzymatic cleavage products that are essentially unavoidable in shells prepared by other *in vivo* protocols and the previously developed *in vitro* method.

As a platform for encapsulating a wide variety of cargo, this method minimizes urea exposure for sensitive biotic cargo, while providing enhanced versatility through the ability to control order of addition for cargo loading of HTP shells. This offers advantages over the combined *in vivo* assembly *+ in vitro* capping protocol, such as the potential to encapsulate bulkier enzymatic cargo or abiotic compounds, like nanoparticles, that are too large to fit through the vacancies in HT wiffle shells and instead require assembly of the entire shell around the cargo. In the case of NrfA, it is a multi-heme enzyme that matures and functions in the periplasm. As such, co-expression with shell proteins is not a suitable method of generating encapsulated NrfA in BMC shells *in vivo*; therefore even the modest encapsulation of 26% of functionally active enzyme is significant as a new-to-nature compartmentalization.

As a powerful tool for bioengineering efforts, our method allows the substitution of native tiles with modified ones, enabling assembly of shells with alternate functionality and properties such as cargo binding affinity and permeability. Peptide extensions known as encapsulation peptides (EPs) are used to pack BMC shells with native enzymes.^34–37^ However, loading shells using EPs is typically inefficient^17,19,38^ and the precise nature of the binding of EPs to shells is not well defined across shell systems. Our method overcomes this shortcoming by using modified tiles, such as _SpyT_BMC-T, thus providing an avenue for covalent cargo encapsulation. *In vivo* encapsulation by targeted loading using affinity tags is time and labor intensive as testing maximum cargo loading can only be done through varying inducer concentrations.^39^ Our *in vitro* method offers more precise control over tile:cargo ratios and allows quicker screening and optimization for shell assembly. Studies have shown that the central pores of the shell tiles determine the permeability of shells to metabolites.^40–42^ Therefore, shell tiles can be designed with mutations that alter overall shell permeability,^43–45^ increasing control of flux of diffusible metabolites.^46^ Control over all the aforementioned parameters and properties by using modified tiles interchangeably increases loading capability and capacity for any specific catalytically relevant cargo.

## Conclusion

Our work presents a new, efficient, and fast method for *in vitro* assembly of bacterial microcompartment shells that can be used in a variety of applications, such as construction of highly specific nanoreactors. This chaotrope-based approach overcomes the barrier to the *in vitro* synthesis of shells presented by insoluble BMC-H sheets that are not viable for shell assembly. The necessary amount of urea needed to disrupt BMC-H sheet structures has proven to be optimal for remarkably robust assembly, allowing increased control and efficiency through the simplicity of combining the three general constituents for shell assembly without additional enzymatic steps. Demonstrated through our ability to incorporate a modified shell tile, our method broadens the scope for bioengineered shells that can be designed for specific applications. The successful encapsulation of biotic and abiotic cargo, both resulting in new-to-nature compartmentalized catalysis, provides evidence of the significant advantage of our method over *in vivo* synthesis and the former *in vitro* shell assembly method.

## Methods

### Expression and purification of BMC shell proteins

BL21(DE3) chemically competent cells (New England Biolabs) were transformed with plasmid DNA containing BMC shell protein sequences according to the vendor’s specifications. Cells were incubated with 50–100 ng of plasmid DNA on ice for 30 minutes and heat shocked at 42°C for 10 seconds. Cells were then placed back on ice for 5 minutes before adding 950 μL of Luria-Bertani broth for 1 hour at 37°C with gentle shaking. 100 μL of recovered cells were plated on kanamycin or ampicillin plates depending on the selective marker in each plasmid and colonies were grown overnight at 37°C.

Colonies of transformed BL21(DE3) cells were grown in 50 mL of Luria-Bertani broth at 37°C overnight. They were grown with 100 μg/mL ampicillin or 50 μg/mL kanamycin, depending on the selective marker in the plasmid. Cells were induced at OD 0.6–0.8 at 600 nm with 50 μL of 1 mg/ml anhydrotetracycline (aTc) (for BMC-T, _SpyT_BMC-T, and BMC-P expressions) or 500 μL of 1 M IPTG (for BMC-H expression) per 1 L culture. The induced cells were incubated at 22°C for 18 hours. The cells were pelleted by centrifugation at 8,000 rpm for 20 minutes at 4°C with a JLA 8.1 rotor in a Beckman Coulter Avanti J-26 XP centrifuge and the supernatant was discarded. Cell pellets were stored at -20°C.

#### Purification of BMC-T, _SpyT_BMC-T, and BMC-P using affinity chromatography

Frozen cell pellets were thawed on ice and resuspended in 30 mL of **Lysis Buffer** (50 mM Tris pH 8, 300 mM NaCl, 10 mM Imidazole, 5% v/v glycerol) with the addition of ½ Sigma protease inhibitor tablet and 200 μL of 2 mg/mL DNase I. Cells were lysed by passage through an Avestin Emulsiflex C3 Homogenizer at 20,000 psi three times. Cell lysate was clarified by centrifugation at 20,000 rpm for one hour at 4°C with a JA 20 rotor in a Beckman Coulter Avanti J-26 XP centrifuge. Supernatants were filtered using a 0.22 μm filter and transferred to clean tubes. Column chromatography was performed using an ÄKTA Start chromatography system (GE Healthcare) and clarified lysates were applied to a 5 mL HisTrap column (GE Healthcare) that was equilibrated with **Buffer A** (20 mM Tris-HCl pH 8, 500 mM NaCl). The column was washed with 6 CV of **Buffer A** then subsequently washed with 6 CV of 98% **Buffer A** and 2% of **Buffer B** (20 mM Tris-HCl pH 8, 500 mM NaCl, 500 mM Imidazole). Protein was eluted over a ten-column volume gradient from 4–100% **Buffer B**. Fractions containing the target protein were identified by SDS-PAGE analysis and were pooled and concentrated with a 3 kDa MWCO Amicon Ultra centrifugal filter (EMD Millipore). Using an ÄKTA Pure protein purification system, proteins were loaded onto the Superdex 200 Increase 10/300 GL by Cytiva. Proteins were eluted in **SEC Buffer** (50 mM Tris-HCl pH 8, 150 mM NaCl) using a flow rate of 0.5 mL/minute. Protein concentrations were quantified by measuring the A280 with a Nanodrop UV-Vis (Thermo) and using the theoretical extinction coefficients (**Table S2**).

#### Purification of BMC-H sheet inclusion bodies

The protocol for purifying BMC-H sheets from inclusion bodies was adopted and slightly modified from Sutter *et al*.^24^ Frozen cell pellets from 1 L culture were resuspended in 30 mL **Hexamer Lysis Buffer** (50 mM Tris-HCl pH 8, 100 mM NaCl, 10 mM MgCl_2_), 200 μL of 2 mg/mL DNase 1, and 100 μL of 10 mg/mL lysozyme. Cells were lysed by 3 passes through an Avestin Emulsiflex C3 Homogenizer at 20,000 psi. 300 μL of Triton X-100 (1% v/v) was added to the lysate and incubated at room temperature with gentle agitation for 20 minutes on an orbital shaker. Insoluble material was separated by centrifugation for 20 minutes at 20,000 rpm with a JA 20 rotor in a Beckman Coulter Avanti J-26 XP centrifuge. The supernatant was discarded and the white pellet was resuspended in 30 mL of **Hexamer Wash Buffer** (50 mM Tris-HCl pH 8, 100 mM NaCl, 10 mM MgCl_2_, 1% v/v Triton X-100) without disturbing the brown cellular debris. The resuspension was transferred to a new centrifuge tube. The centrifugation/wash steps were repeated with 20–30 mL of **Hexamer Wash Buffer** until the pellet was visibly more white and cellular debris was absent. The pellet was then washed once with 30 mL **Hexamer Lysis Buffer** to remove Triton X-100. The pellet was then resuspended in 10 mL of **Hexamer Lysis Buffer** and stored at 4°C.

#### BMC-H sheet denaturation

In an Eppendorf tube, approximately 10–15 mg of BMC-H sheets were spun in a centrifuge at 13,500 rcf for 10 minutes. After decanting the supernatant, the pellet was resuspended in Tris buffer (10mM) with 500 mM urea. The resuspension was incubated at room temperature overnight on an orbital shaker before repeating centrifugation at 13,500 rcf for 10 minutes. The supernatant was transferred to a separate tube without disturbing the remaining pellet. Protein concentration was quantified by measuring the A280 with a Nanodrop UV-Vis (Thermo) and using the theoretical extinction coefficient (**Table S2**).

### Cloning, expression, and purification of cytochrome c NrfA_SpyC_

NrfA_SpyC_ was cloned from an existing vector^35^ containing a codon-optimized sequence for NrfA from *Geobacter lovleyi*, an N-terminal pelB periplasmic localization signal, and a C-terminal Strep-tag II on a pBAD202/D-TOPO backbone. The pBAD backbone was PCR amplified with primers containing sequence overlaps to SpyC. SpyC was PCR amplified with primers that overlapped the pelB region and NrfA. SpyC was inserted by Gibson assembly using the Gibson Assembly Master Mix from New England Biolabs. After mixing PCR products and master mix, samples were incubated for 15 minutes at 50°C. Gibson assembled products were transformed into BL21 *E. coli* and selected for kanamycin resistance. Resistant colonies were picked and plasmid DNA was isolated using New England Biolabs Plasmid DNA Miniprep Kit. Assembly junction sites were then sequenced by Sanger sequencing at the Michigan State University Genomics Core.

NrfA_SpyC_ was expressed in *Shewanella oneidensis* transformed with a pBAD vector containing NrfA with a pelB periplasmic localization sequence, N-terminal SpyCatcher001 and a C-terminal Strep-tag II. Cultures were grown in TB media at 30°C shaking at 160 rpm. Cultures were then induced with a final concentration of 0.02% arabinose at an OD600 between 0.6–0.7 followed by a 16-hour expression at 30°C and 160 rpm and subjected to osmotic shock to lyse the periplasmic space. To perform osmotic shock, cell cultures were pelleted and resuspended in a 20% sucrose solution containing 1 mM EDTA and 30 mM Tris-base at pH 8.0. Cells were then pelleted again and resuspended in ice cold Milli-Q water. The resuspension in ice water was pelleted and the supernatant (lysate) was collected and buffered with **Wash Buffer** (100 mM Tris-HCl, 150 mM NaCl, 0.1 mM EDTA) with addition of one cOmplete-mini EDTA-free protease inhibitor tablet from Sigma Aldrich per liter of culture. Lysate was then filtered and purified with affinity purification on a Strep-Tactin XT 4 flow column from IBA Lifesciences and eluted with 50 mM biotin in **Wash Buffer** at pH 8.0. To further remove contaminants, the eluted protein solution was concentrated on a 10 kDa MWCO centrifugal filter unit from Sigma Aldrich and subjected to size exclusion chromatography on a Superdex 200 Increase 10/300 GL column from Cytiva. All chromatography steps were performed on a Bio-rad NGC chromatography system. Samples were flash frozen with liquid nitrogen and stored at -80°C.

### *In vitro* assembly

To assemble HT and HTP shells, we added BMC-T (and BMC-P) to **IVA buffer** (50 mM Tris-HCl pH 8.0, 150 mM NaCl, 10% v/v glycerol), mixing thoroughly before adding BMC-H, and mixing thoroughly again. The amount of shell proteins added to the reactions was determined by calculating the desired final concentrations of shell proteins. For many of the assemblies reported in the main text, the final concentrations of BMC shell protein tiles were 1 mg/mL BMC-H (16.5 μM tile), 0.33 mg/mL BMC-T (4.8 μM trimer tile), and 1 mg/mL BMC-P (18.3 μM pentamer tile). The assembly reactions were typically incubated for 1 hour to overnight at 4°C.

### SEC purification of assembled shells

An ÄKTA Pure protein purification system was used and samples were loaded onto either the Superdex 200 Increase 10/300 GL by Cytiva or the HiLoad 16/600 Superdex 200 pg by Sigma Aldrich depending upon sample volume. Samples were eluted in **SEC Buffer** using a flow rate of 0.5 mL/minute for the Superdex 200 increase 10/300 or 1 mL/minute for the HiLoad 16/600 Superdex pg.

### Dynamic Light Scattering

Experiments were performed on a Zetasizer Nano S (Malvern). Measurements were taken with 10 acquisitions for 10 seconds each at room temperature using the particle sizing Standard Operating Procedure with default parameters.

### TEM Analysis

10 μL of shell samples were incubated on glow-discharged 300 mesh Formvar-coated TEM grids (Ted Pella) for 1 minute, wicked off with Whatman filter paper, washed 3 times in droplets of water, and stained with 10 μL of 1% uranyl acetate for 1 minute. Excess uranyl acetate was wicked off with Whatman filter paper. Grids were left to air dry and imaged using a Tecnai-12 Transmission Electron Microscope (FEI) operated at 120 kV. Images were recorded on a Gatan 2K x 2K-pixel CCD camera.

### Small Angle X-ray Scattering

Small-angle X-ray scattering (SAXS) measurements were performed at two beamlines: 12-ID-B of Advanced Photon Source (APS) at Argonne National Laboratory and LiX Beamline 16-ID of NSLS2 at Brookhaven National Laboratory. The X-ray energy of APS 12-ID-B beamline was 13.3 keV, and the SAXS data were collected using an Eiger2S 9M detector (Dectris Ltd.). For the LiX beamline, the X-ray energy was 15 keV, and measurements were performed using a Pilatus3 1M detector (Dectris Ltd.). In both cases, flow cells were employed to minimize the potential radiation damage.

### Ru(bpy)_3_ cargo loading and TCSPC photoluminescence spectroscopy

The *in vitro* assembly was performed as outlined above, with the exception that solid Ru(bpy)_3_Cl_2_•6H_2_O (Sigma Aldrich) was dissolved in each of the shell protein solutions to a concentration of ∼13 mM immediately prior to initiating assembly with the addition of BMC-H to the solution of BMC-T and BMC-P. The assembly reaction took place for one hour at room temperature before loading the sample on top of a sucrose cushion (20 mM Tris, pH 7.4, 50 mM NaCl, 30% w/v sucrose) followed by centrifugation overnight at ∼200,000 rcf and 4°C. The next day, the supernatant was discarded, and the pellet was resuspended in HBS (20 mM HEPES, pH 7.4, 50 mM NaCl). The sample was then concentrated and resuspended four times in HBS using 0.5 mL 100 kDa MWCO Amicon Ultra centrifugal filters to filter out trace remaining unencapsulated Ru(bpy)_3_. Protein and ruthenium concentrations were then determined by Bradford assay and inductively coupled plasma atomic emission spectroscopy (ICP-AES), respectively.

Time-correlated single photon counting photoluminescence spectra were measured on a FLS 1000 photoluminescence spectrometer (Edinburgh Instruments) with a PMT-980 detector. An EPL-450 picosecond pulsed diode laser with a 1 second pulse period was used for excitation at 445.8 nm and emission was measured at 620 nm using a 455 nm long pass filter to remove reflected and elastically scattered light from the signal. Decay spectra were fit with either a monoexponential function using Fluoracle software or a biexponential function using OriginLab 2022 software. All samples were measured in 20 mM HEPES, pH 7.4, 50 mM NaCl.

### NrfA cargo loading and methyl viologen activity assay

To encapsulate NrfA_SpyC_ inside HT shells, we mixed the enzyme with _SpyT_BMC-T such that the final ratio of enzymes to assembled shells was approximately 3 enzymes per complete shell (assuming 100% assembly efficiency). The _SpyT_BMC-T+NrfA_SpyC_ mixture was incubated on ice for 1 hour before addition of BMC-H. The assembly reaction mixture was incubated on ice for 1 hour before loading onto a HiLoad 16/600 Superdex 200 pg size exclusion column. The eluted fractions containing assembled NrfA-loaded HT shells were collected and concentrated at 14,000 rcf in a 100 kDa Amicon Ultra centrifugal filter. The presence of NrfA in these fractions was confirmed using UV-Vis spectroscopy, and the concentration was quantified by fitting the heme absorbance peak at 410 nm (**Figure S7**).

We tested the activity of the NrfA inside HT shells using a Michaelis-Menten enzyme activity assay adapted from Campecino *et al*.^28^ with modifications. We added the SEC-purified NrfA-loaded HT shells to **NrfA Activity Buffer** (50 mM Tris-HCl pH 8, 150 mM NaCl, 0.8 mM Methyl Viologen, 0.1 mM Sodium Dithionite) to a target enzyme concentration of 1 nM. This solution is a deep blue color due to the presence of dithionite-reduced methyl viologen. In a 96-well clear-bottom plate mounted on a white light source, we added 190 μL of the enzyme solution to 10 μL of various concentrations of nitrite. Using a Arducam USB-powered RGB camera mounted directly above the plate, we took videos of the wells in the plate at a rate of 1 frame per second. We monitored the progression of the enzyme reaction over time as the solutions in the wells turn from blue to clear. Using MATLAB software, triplicate readings of the kinetic traces were extracted by integrating the pixel intensities within each image’s blue channel corresponding to each well, and the pixel intensities were correlated to the initial reduced methyl viologen concentration of 0.1 mM, as measured by NanoDrop UV-Vis. The rate of change in methyl viologen concentration was found by fitting the first 10 seconds of replicate absorbance readings to a line. The enzyme reaction rates were then fit to a hyperbolic curve using the `curve_fit` function from the `scipy.optimize` Python package to obtain the Michaelis-Menten parameters.

## Acknowledgments

This work was supported as part of the Center for Catalysis in Biomimetic Confinement, an Energy Frontier Research Center funded by the U.S. Department of Energy, Office of Science, Basic Energy Sciences under Award Number DE-SC0023395. The Molecular Foundry at Lawrence Berkeley National Laboratory is supported by the Director, Office of Science, Office of Basic Energy Sciences, US DOE under Contract DE-AC02-05CH11231.

This research used resources of the Advanced Photon Source, a U.S. Department of Energy (DOE) Office of Science user facility at Argonne National Laboratory and is based on research supported by the U.S. DOE Office of Science-Basic Energy Sciences, under Contract No. DE-AC02-06CH11357.

The LiX beamline is part of the Center for BioMolecular Structure (CBMS), which is primarily supported by the National Institutes of Health, National Institute of General Medical Sciences (NIGMS) through a P30 Grant (P30GM133893), and by the DOE Office of Biological and Environmental Research (KP1605010). LiX also received additional support from NIH Grant S10 OD012331. As part of NSLS-II, a national user facility at Brookhaven National Laboratory, work performed at the CBMS is supported in part by the U.S. Department of Energy, Office of Science, Office of Basic Energy Sciences Program under contract number DE-SC0012704.

