## Supporting Information for "A chaotrope-based approach for rapid *in vitro* assembly and loading of bacterial microcompartment shells"

[1]MSU-DOE Plant Research Laboratory, Michigan State University, East Lansing, MI, USA, 48824, [2]Molecular Foundry Division, Lawrence Berkeley National Laboratory, Berkeley, CA, USA, 94720, [3]Cell and Molecular Biology Department, Michigan State University, East Lansing, MI, USA, 48824, [4]Molecular Plant Sciences Program, Michigan State University, East Lansing, MI, USA, 48824, [5]Chemical Sciences and Engineering Division, Argonne National Laboratory, Lemont, IL, USA, 60439, [6]X-ray Science Division, Argonne National Laboratory, Lemont, IL, USA, 60439, [7]Environmental Genomics and Systems Biology Division, Lawrence Berkeley National Laboratory, Berkeley, CA, USA, 94720, [8]Molecular Biophysics and Integrated Bioimaging Division, Lawrence Berkeley National Laboratory, Berkeley, CA, USA, 94720, [9]Department of Biochemistry and Molecular Biology, Michigan State University, East Lansing, MI, USA, 48824.

[†]These authors contributed equally to this work.

Keywords: Bacterial microcompartments, *in vitro*, self-assembly, urea, biotic and abiotic cargo encapsulation, catalysis, confinement

| Protein | Comments | Amino acid sequence |
|---|---|---|
| BMC-H | Wild-type sequence | MADALGMIEVRGFVGMVEAADAMVKAAKVELIGYEK<br>TGGGYVTAVVRGDVAAVKAATEAGQRAAERVGEVV<br>AVHVIPRPHVNVDAALPLGRTPGMDKSA* |
| BMC-T | 6x-Histidine tag tag on N terminus | MHHHHHHMDHAPERFDATPPAGEPDRPALGVLELT<br>SIARGITVADAALKRAPSLLLMSRPVSSGKHLLMMRG<br>QVAEVEESMIAAREIAGAGSGALLDELELPYAHEQLW<br>RFLDAPVVADAWEEDTESVIIVETATVCAAIDSADAAL<br>KTAPVVLRDMRLAIGIAGKAFFTLTGELADVEAAAEV<br>VRERCGARLLELACIARPVDELRGRLFF* |
| $_{SpyT}$BMC-T | 6x-Histidine tag tag on N terminus<br><br>SpyTag-001 flanking Gly-Ser linkers | MHHHHHHMDHAPERFDATPPAGEPDRPALGVLELT<br>SIARGITVADAALKRAPSLLLMSRPVSSGKHLLMMRG<br>QVAEVEESMIAAREIAGAGGGSGGSAHIVMVDAYKP<br>TKGGSGGSGALLDELELPYAHEQLWRFLDAPVVADA<br>WEEDTESVIIVETATVCAAIDSADAALKTAPVVLRDM<br>RLAIGIAGKAFFTLTGELADVEAAAEVVRERCGARLL<br>ELACIARPVDELRGRLFF* |
| BMC-P | 6x-Histidine tag on C terminus | MVLGKVVGTVVASRKEPRIEGLSLLLVRACDPDGTP<br>TGGAVVCADAVGAGVGEVVLYASGSSARQTEVTNN<br>RPVDATIMAIVDLVEMGGDVRFRKDGSSHHHHHH* |
| NrfA$_{SpyC}$ | StrepII tag on C terminus<br><br>SpyCatcher-001 on N terminus | DSATHIKFSKRDEDGKELAGATMELRDSSGKTISTWI<br>SDGQVKDFYLYPGKYTFVETAAPDGYEVATAITFTVN<br>EQGQVTVNGGSGGSAPPKAEQAKIAEIPDGTIDPAV<br>WGKNYPEEYQTWKDTALPTPEGKSKYKKGNDGGK<br>VYDKLSEYPFIALLFNGWGFGIEYNEPRGHVYMMKD<br>QKEIDPSRLKGGGACLTCKTPYAPQLAQKQGVTYFS<br>QSYADAVNQIPKEHQEMGVACIDCHNNKDMGLKISR<br>GFTLVKALDKMGVDQTKLTNQDKRSLVCAQCHVTYT<br>IPKDANMKSQDVFFPWDESKWGKISIENIIKKMRSDK<br>SYGEWTQAVTGFKMAYIRHPEFEMYSNQSVHWMA<br>GVSCADCHMPYTKVGSKKISDHRIMSPLKNDFKGCK<br>QCHSESSEWLKNQVITIQDRAASQYIRSGYALATVAK<br>LFEMTHKQQAAGKQIDQKMYDQAKFYYEEGFYRNL<br>FFGAENSIGFHNPTEAMRILGDATMYAGKADGLLRQ<br>ALTKAGVDVPVKIDLELSKYTNNRGAKKLMFKPEQEL<br>KDPYGPQKWSHPQFEK* |

**Table S1: Amino acid sequences of the proteins used in this study.**

| Protein | Molecular Weight (kDa) | Extinction Coefficient ($M^{-1}$ $cm^{-1}$) | |
|---|---|---|---|
| | | 280 | 410 |
| BMC-H | 10.114 (monomer)<br>60.684 (tile = 6x monomers) | 2980 (monomer)<br>17880 (tile) | |
| BMC-T | 23.085 (monomer)<br>69.255 (tile = 3x monomers) | 12490 (monomer)<br>37470 (tile) | |
| BMC-T-spyTag | 25.026 (monomer)<br>75.078 (tile = 3x monomers) | 13980 (monomer)<br>41940 (tile) | |
| BMC-P | 10.910 (monomer)<br>54.55 (tile = 5x monomers) | 1490 (monomer)<br>7450 (tile) | |
| NrfA-spyCatcher | 62.499 | 96720 | 547000 |

**Table S2: Molecular weights and theoretical extinction coefficients of the proteins used in this study.**

| Assembly Method – Prep # | Molar equivalents of pentamer during assembly/capping | Average # $Ru(bpy)_3$ molecules per shell | $[Ru(bpy)_3]$ in shell lumen (mM) |
|---|---|---|---|
| Combined *in vivo* assembly + *in vitro* capping – Prep 1 | 5x excess | 49 | 5.8 |
| Combined *in vivo* assembly + *in vitro* capping – Prep 2 | 5x excess | 38 | 4.5 |
| One-step *in vitro* assembly – Prep 1 | 1x stoichiometric amount | 20 | 2.3 |
| One-step *in vitro* assembly – Prep 2 | 1x stoichiometric amount | 22 | 2.6 |

**Table S3.** Comparison of $Ru(bpy)_3$ cargo loading experiments by the one-step *in vitro* HTP shell assembly method and the former method combining *in vivo* assembly + *in vitro* capping, Protein concentrations were quantified by Bradford assay and ruthenium concentrations were quantified using inductively coupled plasma atomic emission spectroscopy (ICP-AES). $[Ru(bpy)_3^{2+}]$ in shell lumen value assumes that all protein mass of purified samples represents completely assembled shells with an inner diameter of 30 nm and that the encapsulated $Ru(bpy)_3^{2+}$ molecules are evenly distributed throughout all shells in the sample.

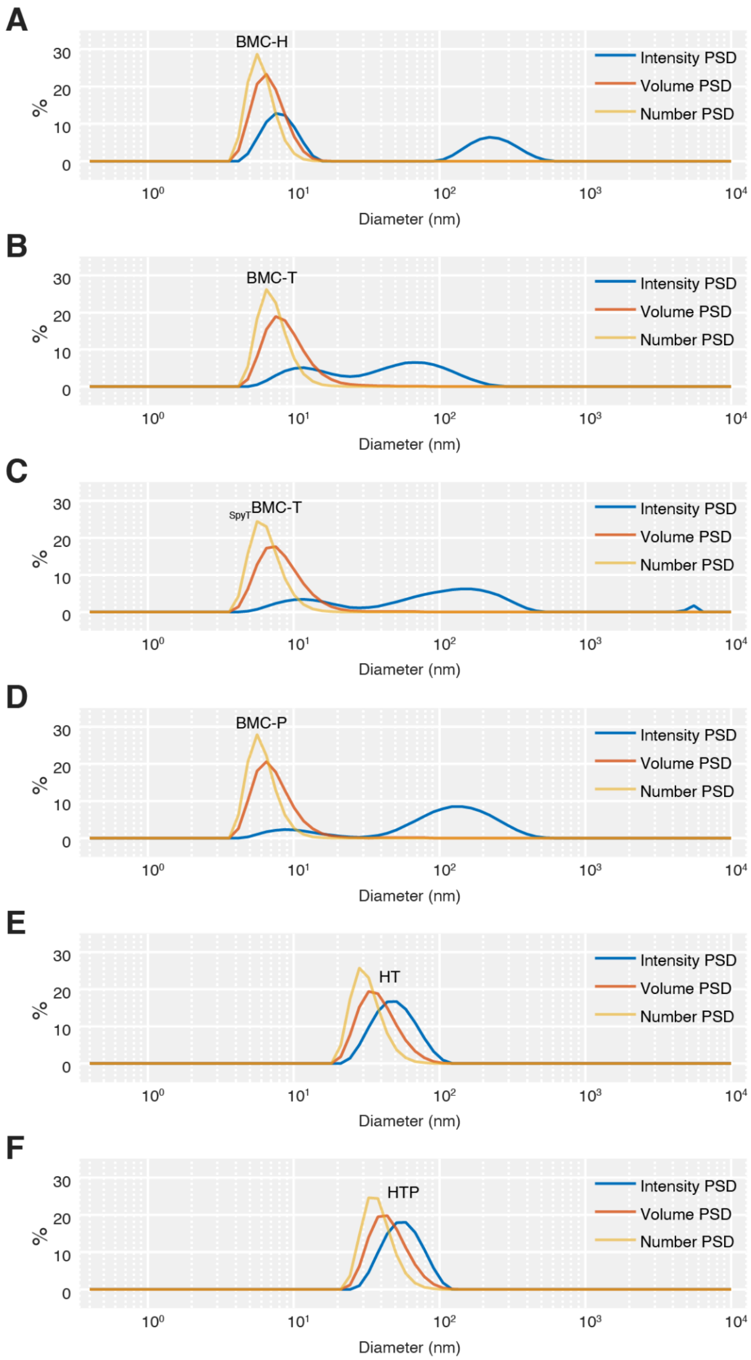


**Figure S1: Typical particle size distributions (PSDs) of BMC shell proteins and their assemblies measured by dynamic light scattering. A.** BMC-H. **B.** BMC-T. **C.** $_{SpyT}$BMC-T. **D.** BMC-P **E.** HT minimal wiffle shells. **F.** HTP minimal shells.

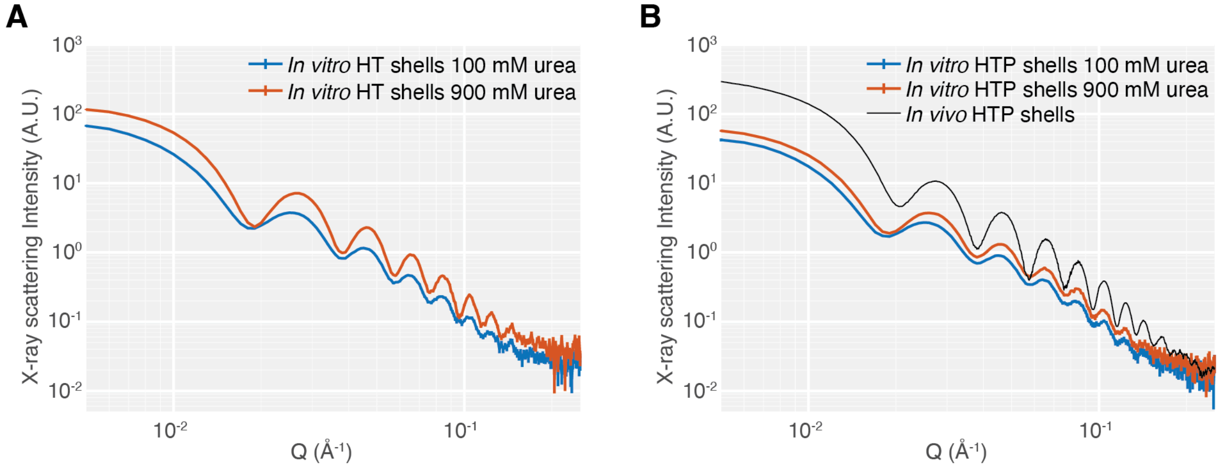


**Figure S2. Experimental small-angle X-ray scattering profiles for HT and HTP shells. A.** HT shells were generated *in vitro* at 100 mM and 900 mM urea. **B.** HTP shells were generated *in vitro* via a one-step assembly at 100 mM and 900 mM urea. For comparison, also shown are *in vivo* HTP shells that were heterologously expressed in *E. coli*. The profiles were offset vertically for clarity.

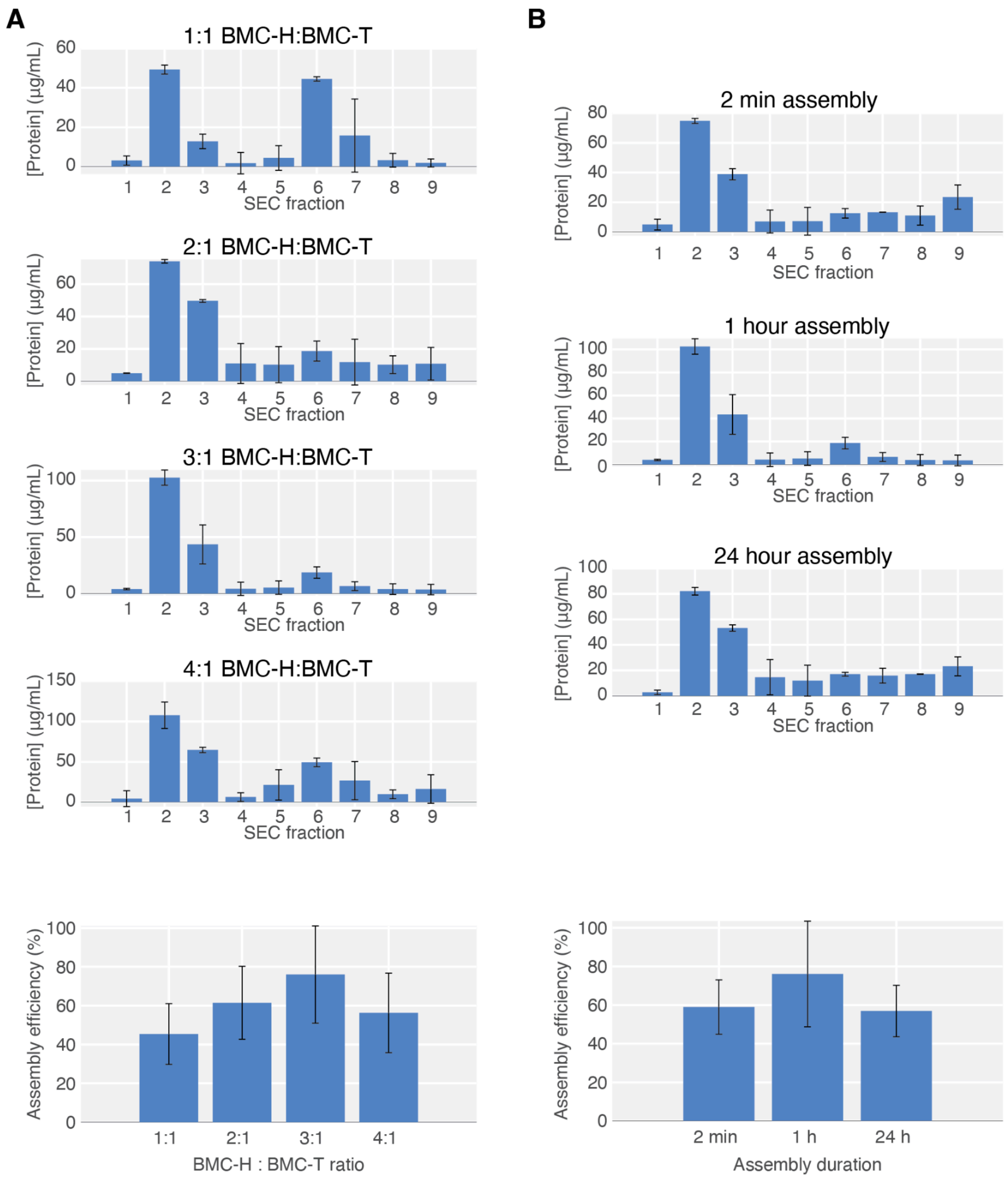


**Figure S3. BCA assay quantification of protein concentrations in SEC elution fractions, and assembly efficiency calculated from BCA-measured protein concentrations. A.** Protein concentrations in SEC elution fractions and assembly efficiency for assemblies in which the ratio of BMC-H:BMC-T was varied. **B.** Protein concentrations in SEC elution fractions and assembly efficiency for assemblies in which the assembly duration was varied.

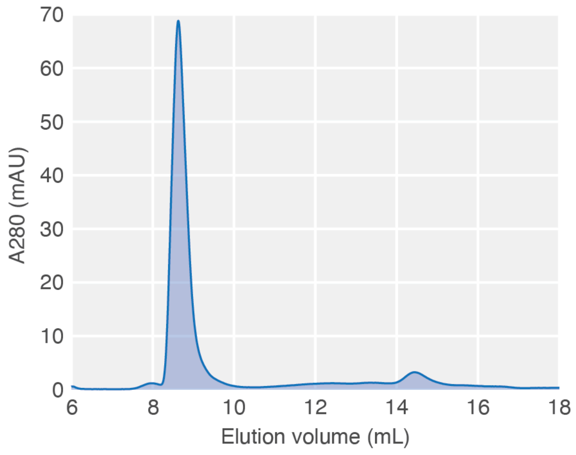


**Figure S4.** SEC chromatogram of a 24-hour HT shell assembly containing 1 mg/mL BMC-H and 0.33 mg/mL $_{SpyT}$BMC-T.

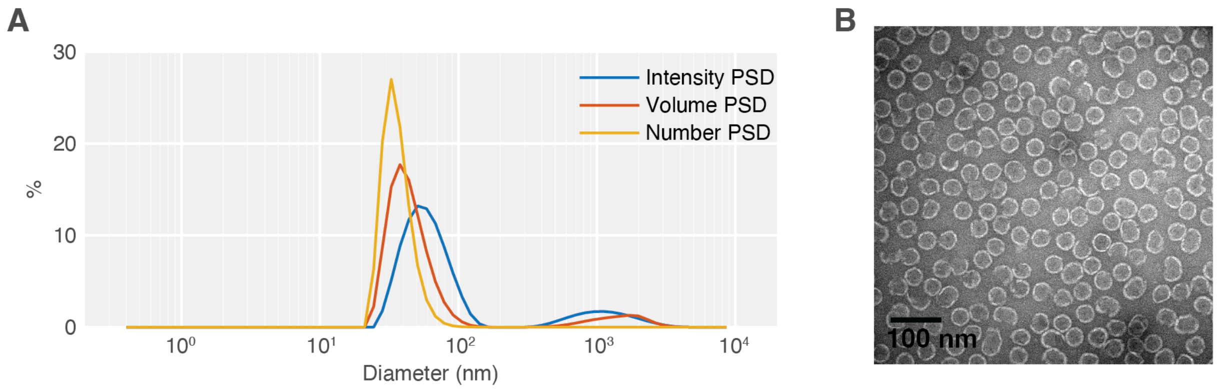


**Figure S5. NrfA-loaded HT shells. A.** DLS particle size distributions of NrfA$_{SpyC}$-loaded HT shells. **B.** TEM micrograph of NrfA$_{SpyC}$-loaded HT shells.

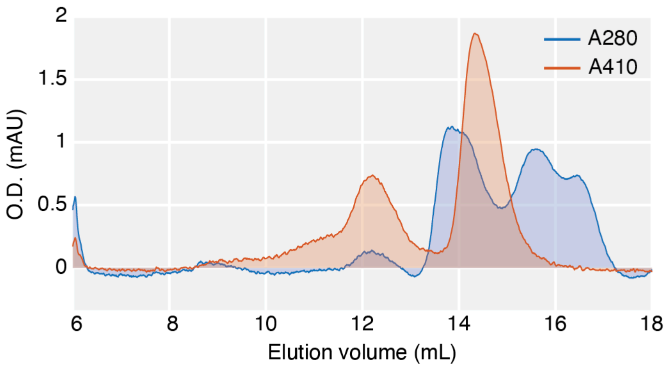


**Figure S6.** SEC chromatogram of a control reaction containing only $_{SpyT}$BMC-T–NrfA$_{SpyC}$ conjugation (no BMC-H added) shows no elution of protein aggregation in the void volume.

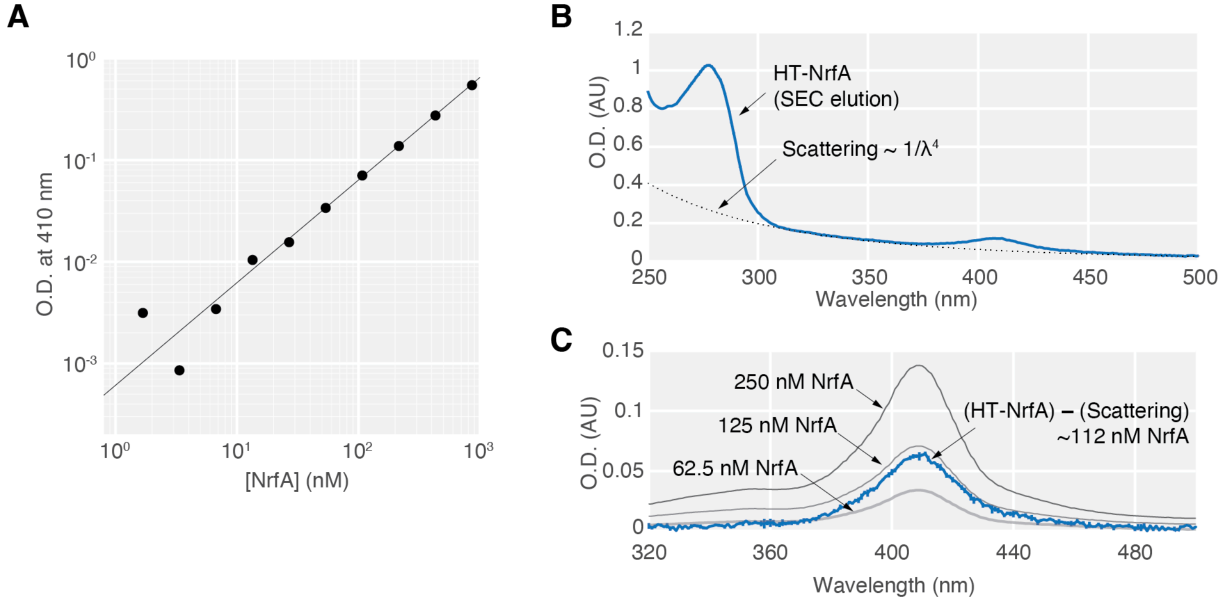


**Figure S7: UV-Vis quantification of NrfA concentration in SEC-purified HT shells. A.** O.D. 410 nm standard curve for serial dilutions of $NrfA_{SpyC}$ stocks with known concentrations. **B.** UV-Vis spectrum for $NrfA_{SpyC}$-loaded HT shells. The dotted line shows a model representing the contribution to the total extinction from scattering. **C.** Enzyme concentration is quantified by isolating the heme Soret absorbance peak at 410 nm. Spectra of several $NrfA_{SpyC}$ solutions with known concentrations are shown for comparison.
